# Soil bacterial biomass and diversity are affected by different furrow-ridge mulched management systems during potato continuous cropping

**DOI:** 10.1101/394551

**Authors:** Weina Zhang, Shuhao Qin, Xuexue Xu, Junlian Zhang, Yuhui Liu

## Abstract

The soil bacterial composition is vital for sustainable agriculture due to its importance in biogeochemical processes in the soil environment. Multiple management systems, such as different furrow-ridge mulched cropping systems, have been established to reduce the damage caused by continuous cropping of potato (*Solanum tuberosum* L.). However, little is known about the responses of soil bacterial biomass and diversity to these systems. In this study, six different ridge-furrow film planting patterns were tested in a 2-year continuous cropping potato field: flat plot without mulch (CK), flat plot with mulch (T1), on-ridge planting with full mulch (T2), on-furrow planting with full mulch (T3), on-ridge planting with half mulch (T4), and on-furrow planting with half mulch (T5). The soil physicochemical properties and bacterial composition were significantly affected by the planting pattern. Mulched soils, especially T2, maintained better soil physicochemical properties than controls. Fully mulched soil maintained higher bacterial biomass and diversity. Among the dominant genera, the abundances of *Nitrosomonadaceae* in T2 and T4 were higher than those in the other treatments. Consequently, compared with the other treatments, on-ridge with mulching patterns resulted in better soil physicochemical properties and high bacterial biomass and diversity, which could reduce the economic losses due to potato production by continuous cropping.

## Introduction

Soil microbial composition, including genetic diversity and quantity, is critical for the maintenance of soil health and quality and directly affects soil function [1]. Many studies have verified that soil microbial communities influence plant growth and productivity, disease resistance, nutrient availability, ecosystem functioning, and plant-soil feedback [2]. Among soil microbes, bacteria play a pivotal role in crop production, supplying nutrients to crops, stimulating plant growth, controlling the activity of plant pathogens, and improving soil structure [3]. For potato planting, the bacteria in the soil are very important for nutrient cycling, decomposition of organic matter, and soil fertility [4]. Concerns have been raised that the alteration of soil microorganisms is one of the major consequences of potato continuous cropping [5, 6]. Therefore, it is important to study the changes in soil bacterial composition and implement effective farming practices to reduce the losses caused by potato continuous cropping.

Potato (*Solanum tuberosum* L.) is an important food crop and plays a crucial role in the agricultural development of an entire region [7]. China is the largest potato producer worldwide and plays an increasing role in the global potato market [8]. However, due to the restriction of land resources, continuous cropping is very common in China [9]. This practice affects soil physicochemical properties, which further result in a series of problems such as increases in pests and diseases, and alterations of the natural microbe balance [10-12]. Therefore, it is important to implement effective practices to reduce the losses from continuous cropping in potato production.

Ridge-furrow mulching planting management systems have been viewed as the simplest and most efficient means for utilizing water. Much research has focused on improving and adjusting farmland moisture [13, 14]. Qiang et al. [15] showed that ridge mulching can prevent soil water evaporation, promote rainfall infiltration, and improve crop root zone soil moisture availability. Planting with mulch can greatly increase soil water storage, utilization rate of soil moisture, and soil physical and chemical properties as well as promote soil microbial breeding [16]. Shi et al. [17] demonstrated that different mulching regimes could improve the activities of catalase, urease, and invertase in the rhizosphere soil of flue-cured tobacco. The increase in catalase activity in the rhizosphere decreased H_2_O_2_-associated damage to plant growth. Changes in urease and invertase activities favored not only the transformation and absorption of soil nutrients at the early growth stage but also the normal maturation and harvesting [17].

Potato soil bacterial diversity cannot be determined using only traditional methods, which often underestimate soil microbial diversity [18]. With the development and adoption of molecular technologies, the Illumina MiSeq second-generation high-throughput sequencing platform, characterized by a high precision of analysis, high sensitivity, and high automation, is extensively used in the medical and microbial fields. Currently, this technique is also successfully applied in studies on microbial diversity [19, 20].

For various crops, previous investigations on the effects of continuous cropping on microbial composition mainly focused on farming practices without mulching. Little is known about soil bacterial diversity under different ridge-furrow mulching planting management systems after potato continuous cropping. Therefore, our study aimed to explore the biomass and diversity of the bacterial community under different management systems using a high-throughput sequencing approach. The information gained will aid in understanding the changes in soil structure caused by potato continuous cropping and will guide in the design of effective measures to increase the production and quality of potato crops.

## Materials and Methods

### Environment of the experimental site

The experiment was carried out at the Experiment Station (104°35’E, 35°33’N) of Dingxi Dry Farming Research Institute of Gansu Agricultural University (Dingxi, China) during 2013-2014. The region is classified as Calcaric Cambisols according to the FAO classification (FAO, 1990), and it has a typical soil of the Loess Plateau. The daily maximum temperatures of Dingxi can reach 38°C in July, whereas minimum temperatures can drop to −22°C in January. The annual average radiation is 5,929 MJ/m^2^, sunshine is 2,477 h per year, and the average long-term annual rainfall is 402 mm (1970-2014) [21]. Bacterial diversity was assessed at the Gansu Key Laboratory of Genetic & Germplasm Enhancement, Gansu Agricultural University (Lanzhou, China).

### Cropping pattern and plant materials

The experiment was performed in a continuous cropping field, and 2014 was the third year of potato continuous cropping. A total of six treatments were used following the method of Qin et al. [21]: two rows of seedlings were planted on a flat plot without any mulch, row space was designed alternating 70 cm with 40 cm (CK); alternating a flat plot mulched with 70 cm plastic film with 40 cm bare land, two rows seedlings with row space of 40 cm were planted in mulched soils (T1); both a wide (70 cm) and a narrow (40 cm) ridge were fully mulched, two rows of seedlings planted on each wide ridge was designed as T2, planted in furrow as T3; alternating a fully mulched ridge (70 cm) with bare land (40 cm), two rows of seedlings planted in the mulched ridges was designed as T4, and planted in bare land as T5.

The black plastic film used in the experiment was 0.01 mm thick. “Xindaping”, a local main cultivar, was planted as the experimental model. The plants were sowed on April 30 and harvested on October 1, 2014; each treatment was repeated three times, and a total of 18 plots were set in a random arrangement. The blocks were designed with a spacing of 2 m, plots were spaced at 1.5 m, and each covered an area of 6.6 × 10 m^2^. The plant row spacing and line spacing were 35 cm and 55 cm, respectively.

### Soil sample collection

The soil samples were collected two weeks before the potatoes were harvested (September 14, 2014) using a dry-sterile brush to brush the surface soil from the potato root into sterile Ziploc bags after removing large particles, broken roots, and stones. The bags were then transferred to the laboratory for analysis. Some samples were placed in an ice box and stored at −80 °C until DNA extraction of soil microbes; the others were air-dried and sieved for determination of the soil physical and chemical properties. The soil was sampled at six points in each plot and then mixed into one sample.

### Microbial culture and determination

The bacteria were cultured in modified peptone-beef extract medium containing 3.0 g beef extract, 5.0 g peptone, 5.0 g NaCl, and 15.0 g agar, with the pH adjusted to 7.0-7.2 [22]. The fungi were cultured in Rose Bengal Medium. The plate count method was used for the determination of microbial quantity.

### Soil bacterial DNA extraction

Soil bacterial DNA was extracted from the three field replicates of six different ridge-furrow film planting patterns using Power Soil DNA Isolation kits (MoBio Laboratories, Inc., US) according to the manufacturer’s recommendations. The quality of extracted DNA was inspected by Gold View staining after 1% agarose gel electrophoresis (AGE).

### PCR amplification and MiSeq sequencing analysis

Amplifications of bacterial 16S rRNA genes were performed with an ABI GeneAmp 9700 PCR instrument with specific synthetic primers: 515F 5′-GTGCCAGCMGCCGCGG-3′ and 907R 5′-CCGTCAATTCMTTTR-AGTTT-3′. The PCR reaction was carried out with a total reaction volume of 20 µl, containing 4 µl 5×FastPfu Buffer, 2 µl 2.5 mM dNTPs, 0.8 µl Forward Primer (5 µM), 0.8 ml Reverse Primer (5 µM), 0.4 µl FastPfu Polymerase, and 10 ng purified soil DNA as the template, with ddH_2_O added. The PCR program consisted of an initial denaturation step of 95°C for 3 min, followed by 27 cycles at 95°C for 30 s, 55°C for 30 s, and 72°C for 45 s and a final extension step at 72°C for 10 min. The triplicate PCR products were assessed with 2% agarose gel electrophoresis, purified using an AxyPrepDNA gel recovery kit (Axygen Inc., Union City, CA, USA), and quantified after the product was eluted with Tris-HCl. High-throughput sequencing was performed using Illumina MiSeq platforms at the Gansu Key Laboratory of Crop Genetic and Germplasm Enhancement.

### Statistical analysis

The 16S rDNA sequences of the six treatments were optimized on the basis of the overlapping relationship between PE reads, and then, pair-reads were merged as a single sequence. Operational taxonomic units (OTUs) were defined by a 3% difference, and each OTU represented a type sequence [23]. The most similar bacterial species were found in GenBank using BLAST searches. Bacterial diversity was assessed using species richness, characterized as the number of species (https://www.mothur.org/wiki/Chao); the Shannon-Wiener diversity index, defined as species diversity in a community (https://www.mothur.org/wiki/Shannon); and sequencing depth, measured with Mothur. Moreover, principal component analysis (PCA) was performed using CANOCO software for Windows 4.5 to establish possible relationships among soil bacterial community distribution for the different planting patterns.

Statistical analyses of the data were performed using Origin 8.0 (Microcal Software, Northampton, MA, USA), SPSS statistical software version 10.0, and the Student’s t-test (P < 0.05 was considered significant).

## RESULTS

### Effects of planting patterns on soil physicochemical properties

The physicochemical properties of the soil profiles for the six different planting patterns are summarized in Table 1. High levels of relative water content were detected for on-ridge and on-furrow planting with mulching (T2, T3, T4, and T5). Compared to the other treatments, a more moderate pH value and a lower electrical conductivity (EC) value were observed for T3 and T2, respectively. Among all six planting patterns, T2 had the highest available nitrogen (N) content; CK, T3, and T5 had the highest available potassium (K) levels; and T3 had the highest available phosphorus (P) level. These results indicated that the physicochemical properties were significantly affected by the different planting patterns, among which the mulched patterns performed better than that the control.

**Table 1.** Physicochemical properties of soil profile

| Treatment | Relative<br>water<br>content /% | pH | Electrical<br>conductivity<br>(EC)<br>($\mu\text{S}/\text{cm}$) | Available<br>nitrogen<br>($\text{mg}\cdot\text{kg}^{-1}$) | Available<br>potassium<br>($\text{mg}\cdot\text{kg}^{-1}$) | Available<br>phosphorus<br>($\text{mg}\cdot\text{kg}^{-1}$) |
| --- | --- | --- | --- | --- | --- | --- |
| CK | 7.26b | 7.74a | 284ab | 15.0ab | 236a | 26.0b |
| T1 | 8.90ab | 7.64ab | 276ab | 12.4c | 124b | 29.2ab |
| T2 | 11.04a | 7.45ab | 232a | 16.6a | 112b | 38.7ab |
| T3 | 10.51a | 7.29b | 241a | 12.1c | 137ab | 40.4a |
| T4 | 10.25a | 7.47ab | 244a | 14.7b | 107b | 26.6ab |
| T5 | 10.54a | 7.56ab | 283ab | 16.5a | 152ab | 36.5ab |

### Effects of planting patterns on soil bacterial biomass

The soil bacterial biomass results at three time points for the six investigated treatments are shown in Table 2. Soil bacterial numbers varied from 42.9 to 74.5, 74.6 to 138.5, and 54 to 109.7 × 10^5^ CFU·g^−1^ on April 13, July 8, and September 15, respectively. A higher level of soil bacterial biomass was found on July 8 among the three time points. For all treatments, soil bacterial numbers ranked in the order of T2 > T4 > T5 > T3 > T1 > CK on Apr 13, T2 > T3 > T4 > T5 > CK > T1 on July 8, and T2 > T5 > T4 > T3 > T1 > CK on September 15. The results revealed that the number of bacteria first increased and then decreased, and the highest levels were found for T2 at all three time points. These results indicated that on-ridge planting with full mulch is beneficial for soil bacterial biomass.

**Table 2.** Effect of soil bacterial number under different ridge-furrow film continuous planting patterns

| Treatment | Bacterial number ($\times 10^5 \text{CFU} \cdot \text{g}^{-1}$) | | |
| --- | --- | --- | --- |
|  | Apr-13 | Jul-8 | Sep-15 |
| CK | 42.9c | 88.4de | 54.0c |
| T1 | 57.4b | 74.6e | 67.1bc |
| T2 | 74.5a | 138.5a | 109.7a |
| T3 | 58.2b | 127.0ab | 74.7b |
| T4 | 64.3b | 112.3bc | 82.1b |
| T5 | 62.2b | 98.9cd | 82.8b |

### 16S rDNA optimizing sequence statistics

Following quality and length filtration, 183,389 reads of 16S rDNA were obtained, with a total length of 72,683,859 bp and an average length of 396.34 bp. From each sample, the number of obtained optimized sequences varied from 9,876 to 20,291 and ranked in the order of T1 > T2 > T5 > CK > T4 > T3.

### Soil bacterial diversity indices and richness

The bacterial species richness and diversity results for the six different planting patterns are shown in Table 3. More than 90% of the coverage indices indicated that the true bacterial diversity in the tested samples was reflected by the sequencing results at a similarity level of 0.03. The Chao value indicated the richness of the bacterial diversity in soil. In the early stage (8-July), the bacterial richness was not affected by the planting patterns (data not shown). However, at a later growth stage (15-September), the bacterial richness in all tested soils was higher than that in CK. Two fully mulched planting patterns had higher bacterial richness values than those of the other planting patterns, which ranked as T3 > T2 > T1 > T5 > T4 > CK. In addition, the diversity of the microbial community from each sample was estimated using the Shannon-Wiener index. In the early stage, the bacterial diversity from all tested samples was higher than that of the control (data not shown). At a later growth stage, the highest bacterial diversity was also observed for fully mulched planting patterns (T2 and T3) among all samples (T2 > T3 > T5 > T1 > T4 > CK). Thus, the bacterial richness and diversity were significantly affected by planting patterns, and two fully mulched planting patterns (T2 and T3) had the highest bacteria diversity.

**Table 3.**
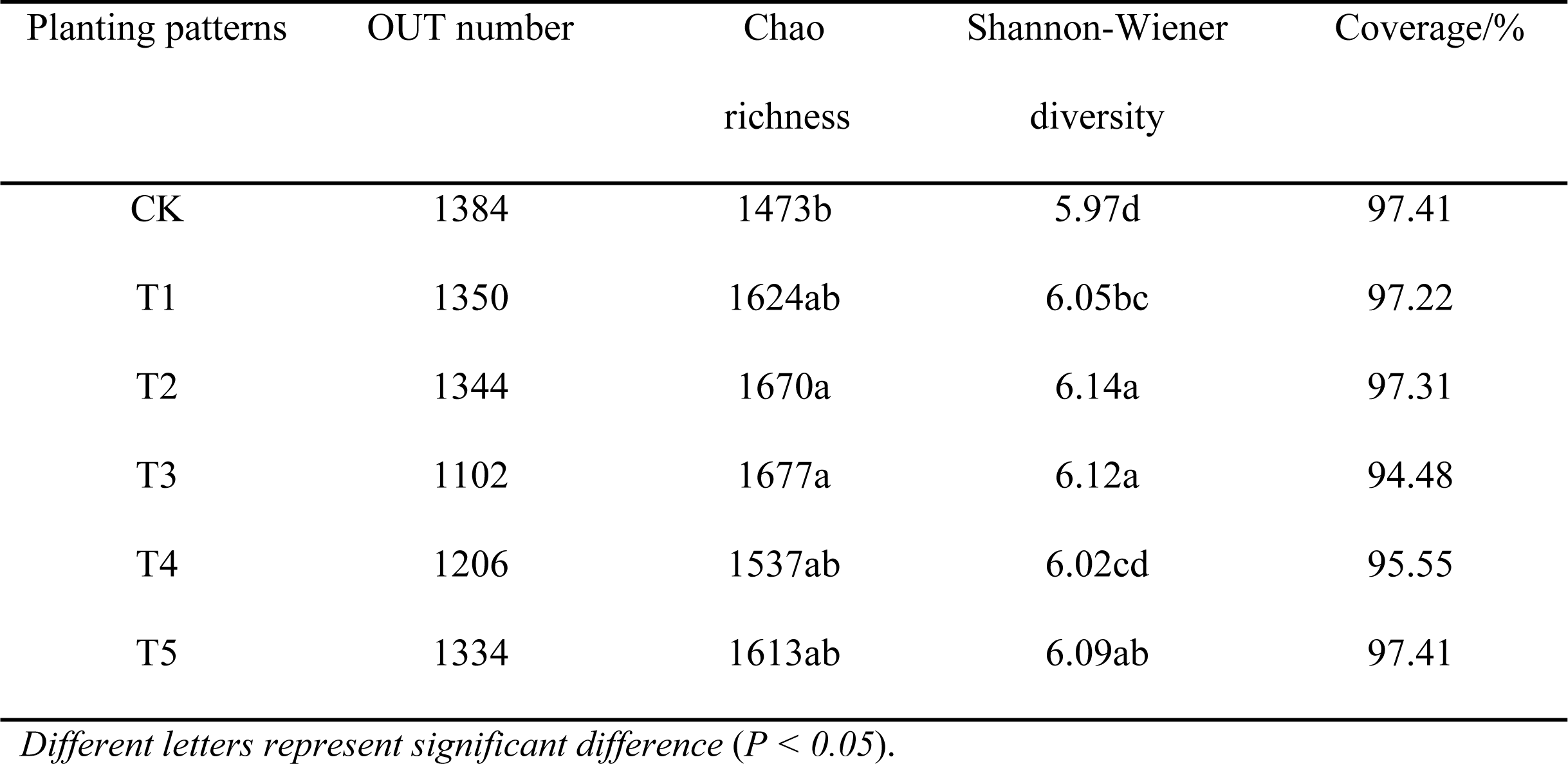
Diversity and richness of soil bacterial community in different furrow-ridge mulching planting patterns (15-September)

### Bacterial community composition in soil

The bacteria were classified into 33 phyla, 73 classes, 163 orders, and 439 genera. At the phylum level, bacteria in continuous cropping soil mainly included *Acidobacteria, Actinobacteria, Bacteroidetes, Proteobacteria, Planctomycetes, Gemmatimonadetes, Verrucomicrobia, Chloroflexi*, and *Firmicutes* (Figure 1). Among them, *Acidobacteria* and *Proteobacteria* were the dominant communities, followed by *Bacteroidetes* and *Planctomycetes*. The taxonomic composition of *Acidobacteria* in the six treatments varied from 26.0 to 31.4%, in the order of T4 > CK > T3 > T2 > T1 > T5. *Proteobacteria* varied from 23.2 to 34.5%, in the order of T5 > CK > T4 > T3 > T2 > T1. The proportion of *Bacteroidetes* varied from 11.4 to 16.0%, whereas that of *Planctomycetes* varied between 7.8 and 9.2%.

**Figure 1.**
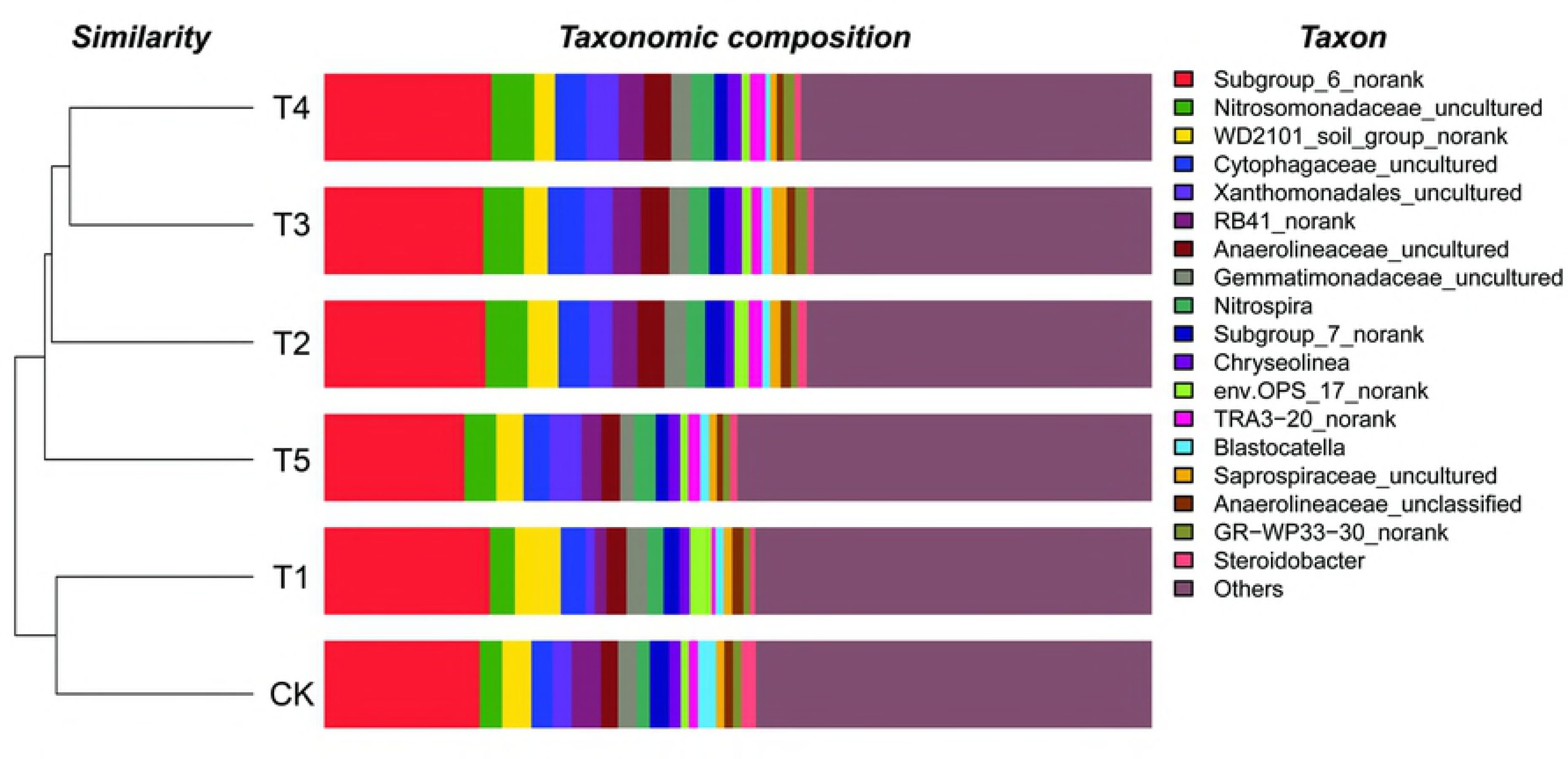
Bacterial phyla in the soils from the different planting patterns.

The color gradients and similarity degrees in the heatmap shown in Figure 2 indicate the diversity and richness of the bacterial communities of the six tested soils. The bacteria in continuous cropping soils were classified into 439 genera, including *Subgroup_6_norank, Blastocatella, Bryobacter, Chryseolinea, Cytophagaceae, Gemmatimonadaceae, Steroidobacter*, and *Xanthomonadales*. *Subgroup_6_norank* in the six tested soils varied from 14.0% to 20.2%. Among the different planting patterns, the abundances of the genera for most of the mulched soils (T1, T2, T3, and T4, accounting for 19.2 to 20.22%) were higher than that in the control (CK, 18.77%). Other dominant genera were *RB41_norank, WD2101_soil_group_norank, Nitrosomonadaceae, Cytophagaceae, Xanthomonadales, Gemmatimonadaceae, Anaerolineaceae*, and *Nitrospira*, and the average abundance of the six samples changed from 2.2% (*Nitrospira*) to 4.12% (*Nitrosomonadaceae_uncultured*). The abundances of *Nitrosomonadaceae_uncultured, Cytophagaceae_uncultured, Xanthomonadales_uncultured*, and *Anaerolineaceae_uncultured* were higher for the on-ridge and on-furrow planting patterns (T2, T3, T4, and T5) than for the others (CK and T1). In addition, the abundances of *Nitrosomonadaceae_uncultured* for the ridge planting pattern (T2 and T4, accounting for 5.09 and 5.09%) were higher than those for the furrow (T3 and T5, accounting for 4.93 and 3.83%) and other planting patterns (CK and T1, accounting for 2.72 and 3%).

**Figure 2.**
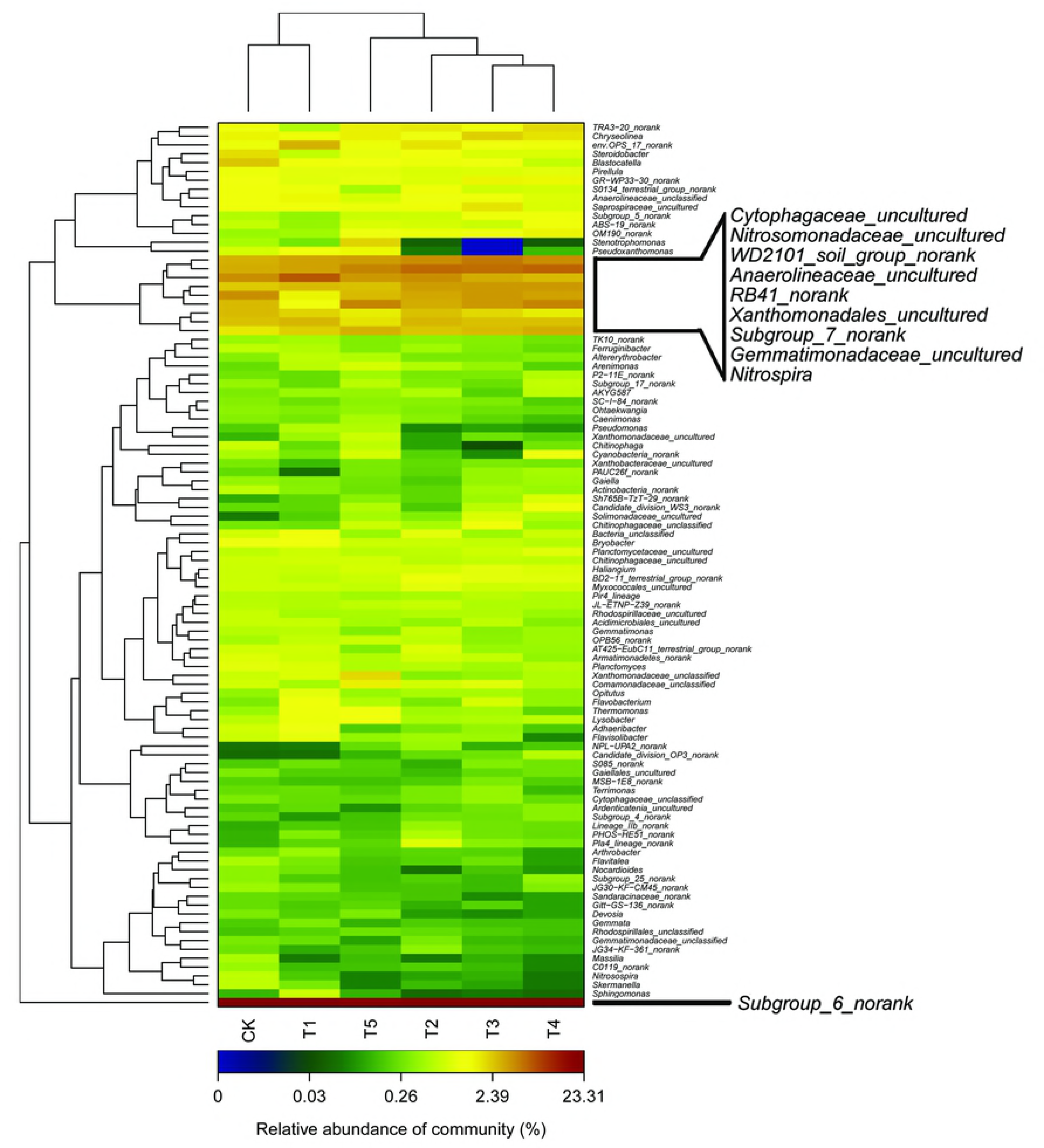
Heatmap of soil bacterial genera in the different soils. Based on the vertical and horizontal clustering, the results are presented as operational taxonomic units (OTUs) in the figure. A detailed statistical analysis at the OTU level, representing bacterial species, revealed a complex community structure of the rhizosphere. The similarities and differences in community composition from different samples under the classification level are reflected by the color gradient and similar degrees.

Cluster analysis at the genera level revealed that the bacterial compositions in the soils from the different treatments could be classified into two groups: CK and T1 and T2, T3, T4, and T5. This indicates obvious differences in bacterial diversity and richness between mulched ridge cropping and flat cropping.

### PCA of the soil bacterial community in different furrow-ridge mulching planting patterns

The first three (PC1, PC2, and PC3) components accounted for 34.85%, 19.14%, and 12.89% with a whole variance of 66.88% (Figure 3). The large interval distribution of the data points between different treatments indicates the obvious effects of the different management systems on the composition of the bacterial community in the soil. Therefore, the PCA results confirmed that different furrow-ridge mulching planting patterns can change the soil quality, resulting in variations in the soil bacterial profile (Figure 3).

**Figure 3.**
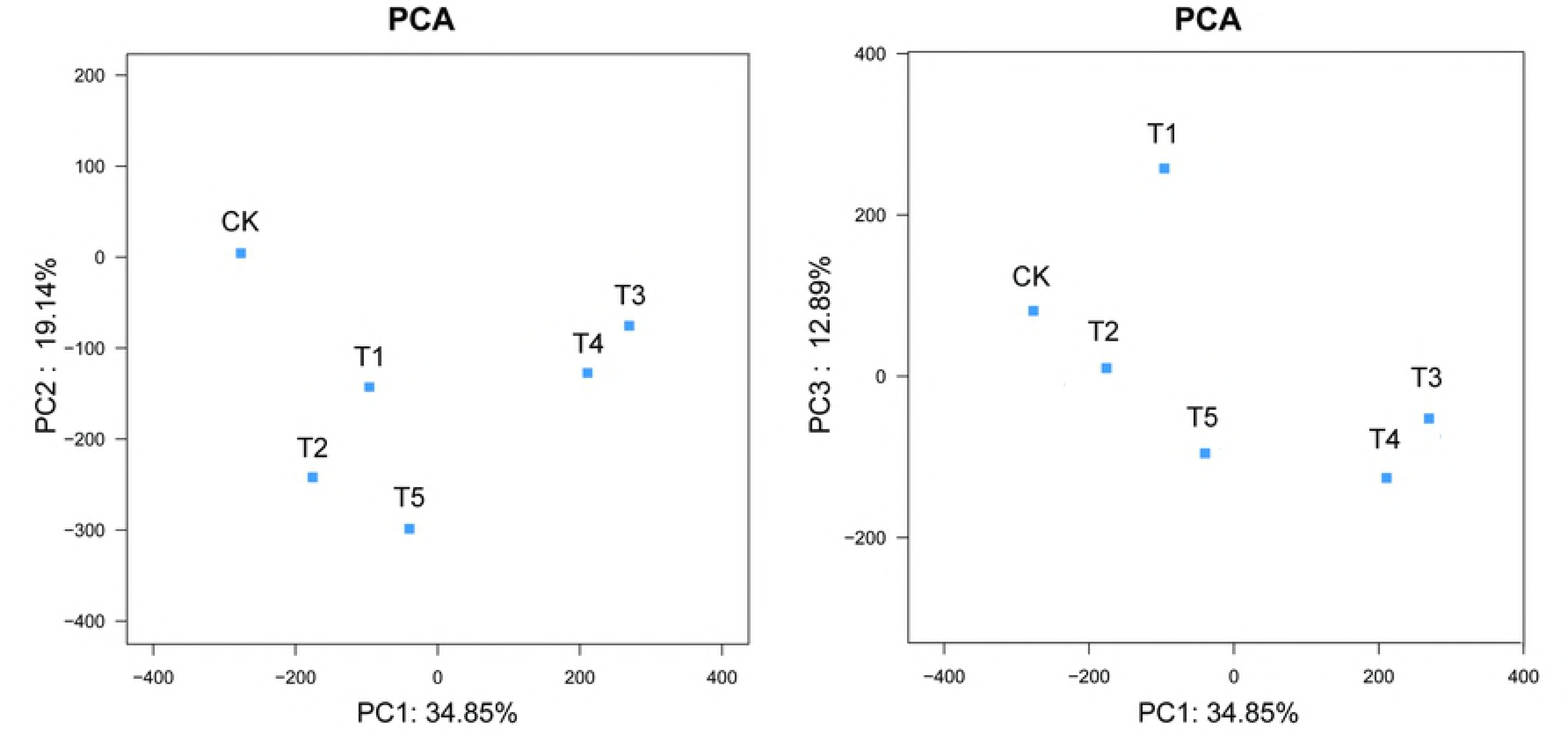
Principal component analysis (PCA) of the soil bacterial community in the different soils.

### Yield of potato

Compared with the control (CK), all five mulched cropping systems exhibited a significant increase in potato yield in both 2013 and 2014. The yield could be ranked as T1 > T2 > T3 > T4 > T5 > CK in 2013 and T2 > T1 > T4 > T5 > T3 > CK in 2014. The yield from CK decreased in 2014 compared with 2013; however, other treatments showed no change or obvious trends of increase.

## Discussion

Continuous cropping is very common and has drawn public attention in China [24]. It is imperative to understand the mechanism of the damage caused by continuous cropping and take measures to reduce the losses that result from it. The mechanism of the damage caused by crop continuous cropping is very complicated, although an imbalance in the structure of the soil microbial population is the main factor [25]. Changing the biomass of soil bacteria and the structure of the microbial community can negatively affect microbial function, leading to decreases in soil nutrients and fertility [26, 27]. Our results showed that both bacterial biomass and composition were markedly affected by mulching during the process of potato continuous cropping.

Over a long period of continuous cropping, the number of soil bacteria and actinomycetes sharply decrease, whereas that of fungi increases [28, 29]. Soil fertility can change from “bacterial” to “fungal” conditions. “Fungal” soil causes more serious soil diseases, mainly because of the enrichment of certain microbial populations, especially plant pathogenic fungi, which are not conducive to the soil microbial population balance but are conducive to plant root diseases [30]. This type of change in soil conditions might be an important factor responsible for the loss of potato production and quality [31]. Our results indicated that a high level of bacterial biomass in the soil was associated with fully mulched practices, especially full mulch with an on-ridge pattern (T2) (Table 2 and Table 3). Interestingly, our results also revealed a lower level of fungal biomass in most potato growth periods from fully mulched soils (Table 4). Thus, the decrease in bacterial biomass and increase in fungal biomass could possibly be directly or indirectly prevented by using mulching management. Moreover, we also verified that the soil from T2 maintained better soil fertility, such as a higher relative water content, moderate pH value, lower electrical conductivity, and higher available nitrogen and phosphorus (Table 1). Wang et al. [32] reported similar results regarding the effects of the plastic film cultivation pattern on soil fertility. Therefore, we conclude that fully mulched practices help maintain a higher bacterial biomass and soil fertility during the process of continuous cropping.

**Table 4.** Effect of soil fungal number under different ridge-furrow film continuous planting patterns

| Treatment | Numbers of fungus ($\times 10^2 \text{CFU} \cdot \text{g}^{-1}$) | | |
| --- | --- | --- | --- |
|  | Apr-13 | Jul-8 | Sep-15 |
| CK | 90.1c | 117.8a | 63.6b |
| T1 | 79.1c | 43.8cd | 69.2ab |
| T2 | 17.8e | 54.1c | 73.8a |
| T3 | 43.6d | 38.7d | 62b |
| T4 | 111.5b | 48.4cd | 59.9b |
| T5 | 145.5a | 66.3b | 66.4ab |

Farming management practices have been shown to affect the structure of the microbial community [33]. In our previous study, we found that different furrow-ridge mulching management systems significantly affected soil fungal diversity [21]. Similarly, different systems also markedly affected bacterial diversity, although common dominant bacteria were found in the different mulched soils both at the phylum and genus levels (Figures 1, 2, and 3). The composition of dominant bacteria at the phylum level was consistent with previous investigations [19; 34, 35], further indicating the high reliability of the data from this study. *Nitrosomonadaceae*, a dominant genus in continuous potato cropping soil, comprise a monophyletic phylogenetic group within *betaproteobacteria* [36, 37]. All these cultivated representatives are lithoautotrophic ammonia oxidizers and are crucial for soil function [38, 39, 40]. Interestingly, the richness of *Nitrosomonadaceae* in soil from on-ridge planting with full or half mulch (T2 and T4, accounting for 5.09 and 5.09%) was much higher than that in the control treatment (CK). *Nitrosomonadaceae* plays major roles in controlling the nitrogen cycle in terrestrial, freshwater, and marine environments and in wastewater treatment processes [36, 41, 42]. This also explains why the soil from the on-ridge planting with mulch system had higher available nitrogen content (Table 1). Additionally, although the functions of most of the other genera regarding soil nutrients and fertility are currently not clear, our results from the PCA confirmed the clear effect of different furrow-ridge mulching management systems on diversity at the genus level (Figure 3). Furthermore, we have confirmed higher potato production from both furrow and ridge mulched soils (Figure. 4). In summary, these results indicate that the beneficial effects on bacterial diversity from mulched soils also benefit soil fertility and potato production during the process of potato continuous cropping.

**Figure 4.**
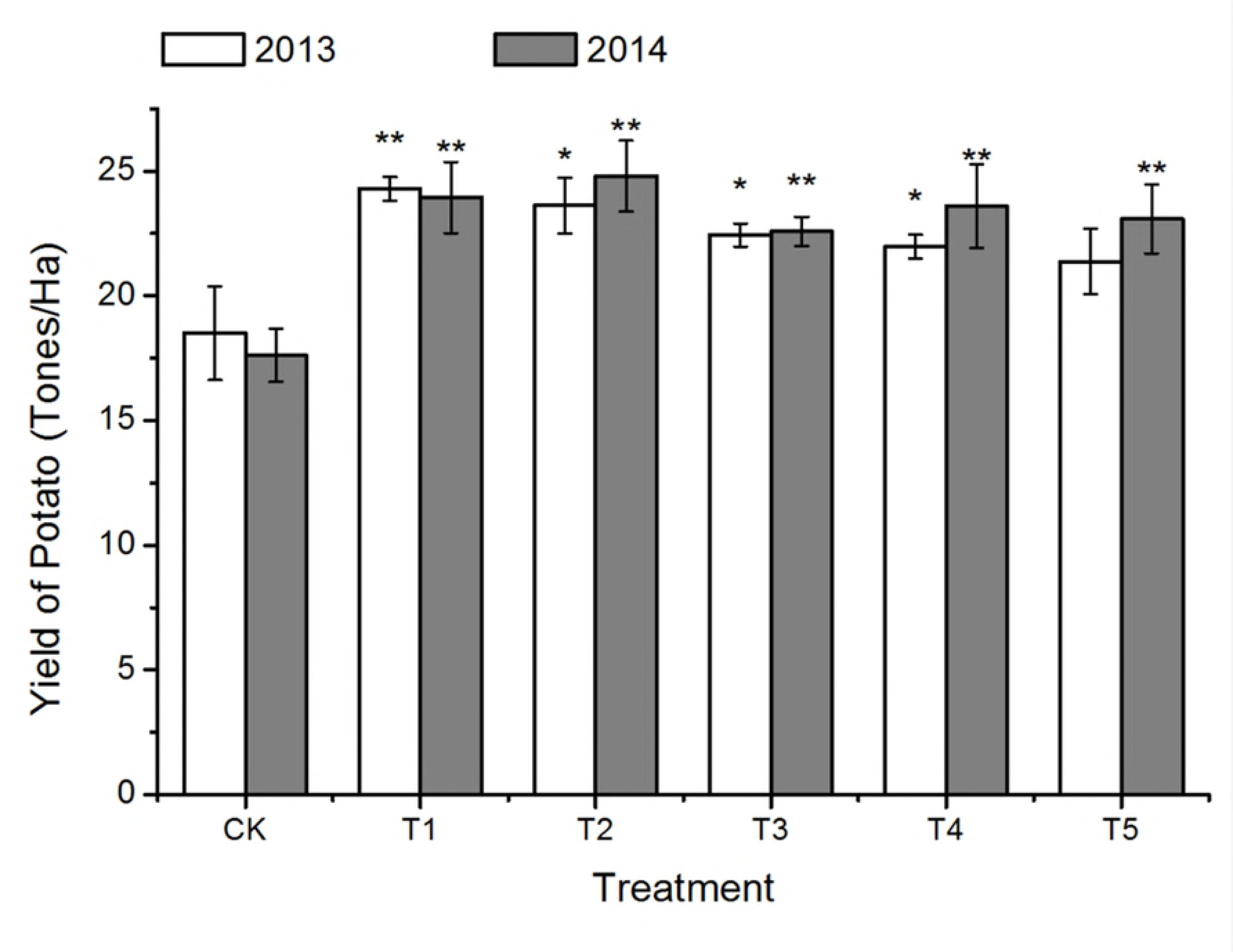
Yield of potato from different treatments. Asterisks indicate a significant difference from the control in the same year at P < 0.05 (*) and P < 0.01 (**).

In conclusion, different furrow-ridge mulching management systems clearly affected soil physicochemical properties, bacterial biomass, and bacterial diversity during the process of potato continuous cropping. Mulched soil, especially on-ridge planting with full mulch (T2), maintained better soil fertility and higher bacterial biomass than controls. A higher proportion of *Nitrosomonadaceae* was found in furrow-ridge mulched soils than in the control group. These changes affected both the soil function and reduced the loss of potato production due to the effects of continuous cropping.

## Author Contributions

SQ and WZ conceived the study and participated in its design and prepared the manuscript. WZ and XX performed the experiments and collected, analyzed, and deposited the data. JZ and YL edited the final draft and revised the manuscript. All authors have read and approved the manuscript.

## Acknowledgments

We sincerely thank all the students and staff of Dingxi Rainfed Agricultural Research Institute for their assistance with the field work.

## Ethical statement

This article does not contain any studies with human participants performed by any of the authors.

